# BIC Defacing Algorithm

**DOI:** 10.1101/275453

**Authors:** Vladimir S. Fonov, Louis D. Collins

## Abstract

Public distribution of imaging information from several MRI data processing projects at the BIC has lead to the development of this DEFACING algorithm that is used as part of anonymisation process. Key features of the algorithm include: the defacing should modify voxels associated with face of the subject, making a rendering of the face unrecognisable, it should work on data in the native scanner space and it should not significantly affect subsequent data processing outcome (model based registration, brain extraction, brain tissue classification and brain segmentation). The algorithm is implemented using the MINC library and source code is publicly available. The effect of the defacing algorithm on the data processing was verified using the ICBM database of 152 scans of young adults.

## 1 Introduction

### 1.1 Background

Current NIH regulations for data sharing [1] have the requirement “The rights and privacy of people who participate in NIH-sponsored research must be protected at all times.”. Currently there are a number of research projects that are making MRI data publicly available, including but not limited to the NIH pediatric database project [2], the Biomedical Informatics Research Network [3] and the Alzheimer’s Disease Neuroimaging Initiative [4]. Many of these projects rely upon a fairly standardised initial data processing pipeline [5], [2].

Several somewhat rudimentary techniques have been developed to address the problem of data anonymisation, for example the BIRN method [6] and John-Doe algorithm [7], although providing reasonable deidentification of the facial features these methods are not suitable for anonymisation of the native images in the scanner space. The main cause of this is that they alter the data significantly and thus affect subsequent processing. The algorithm presented in this paper is designed in a way that makes it possible to deface data in the native space without affecting the result of the following data processing.

The algorithm is evaluated using a quite common imaging protocol consisting of 3 anatomical scans (T1 weighted, T2 weighted and PD weighted) but could be easily adapted to different protocols. Initial data processing consists of the following steps:

1. B0 Intensity inhomogeneity correction using N3 [8]
2. Linear intensity normalization to correct for global intensity fluctuations
3. Linear registration of all 3 modalities into MNI-space, using the ICBM-152 model [9]
4. Brain Extraction using the Brain Extraction Tool (BET) [10]

### 1.2 Overview of existing defacing techniques

Other groups have successfully developed several MRI de-identification techniques:

- The BIRN method [6] simply assigns a zero value to the voxels in the facial part of the mri scan. Whilst this approach can guarantee removal of the chosen facial features, the nature of the operation will often seriously alter the results of subsequent automated processing, most notably registration to MNI-space. As such, this method is typically applied to images after stereotaxic registration.
- The John-Doe algorithm [7] smoothes the voxel values in the facial area and thus removes some recognisable features.

Both of these methods lack verification or quantification of the effect they have on subsequent automatic data processing steps (tissue classification for example).

### 1.3 Requirements of a defacing procedure

A successful defacing procedure should satify all of the following constraints. First, it should make identification of a subject by surface rendering of the face impossible. It must be developed in such a way that it can be applied to the data in the native scanner space to enable comparison of different post-processing techniques when raw data is released. It should not affect any data processing that follows to such an extent that it changes the eventual results in a statistically measurable fashion. Any changes to the subjects face should be limited to the soft tissue structures and not modify the skull such that rigid body skull based registration methods are not compromised.

## 2 Methods

### 2.1 ICBM database

Within the ICBM project, MRI data from 152 young normal adults (18–43.5 years) were acquired on a Philips 1.5T Gyroscan (Best, Netherlands) scanner at the Montreal Neurological Institute [11]. The T1w data were acquired with a spoiled gradient echo sequence (sagittal acquisition, 140 contiguous 1-mm thick slices, TR = 18 ms, TE = 10 ms, flip angle 30°, rectangular FOV of 256 mm SI and 204 mm AP). The T2w/PDw data were acquired as a dual contrast fast spin echo sequence acquired in the axial direction with TR = 3300 ms, TE1 = 34 ms, TE2 = 120 ms, FOV of 256 mm AP and 224 mm LR, with a 2 mm slice thickness. The Ethics Committee of the Montreal Neurological Institute approved the study, and informed consent was obtained from all participants.

### 2.2 Defacing algorithm

The initial data processing is performed on the raw T1, T2 and PD data. The following information is then used in the defacing algorithm: Registration parameters for T1, T2 and PD modalities from scanner native space to stereotactic space [9]: *R*_*t*1_, *R*_*t*2_, *R*_*pd*_ and brain mask [10]: *M*_*b*_.

The area subjected to the deformations (face) has to be specified in advance in the form of the binary face mask in the stereotactic space *M* _*f*_, it is combined with subject’s brain mask: *M*_*s*_ *= M*_*f*_ \ *M*_*b*_ to ensure that the deformations will not affect the brain tissues, see Figure 1

**Figure 1.**
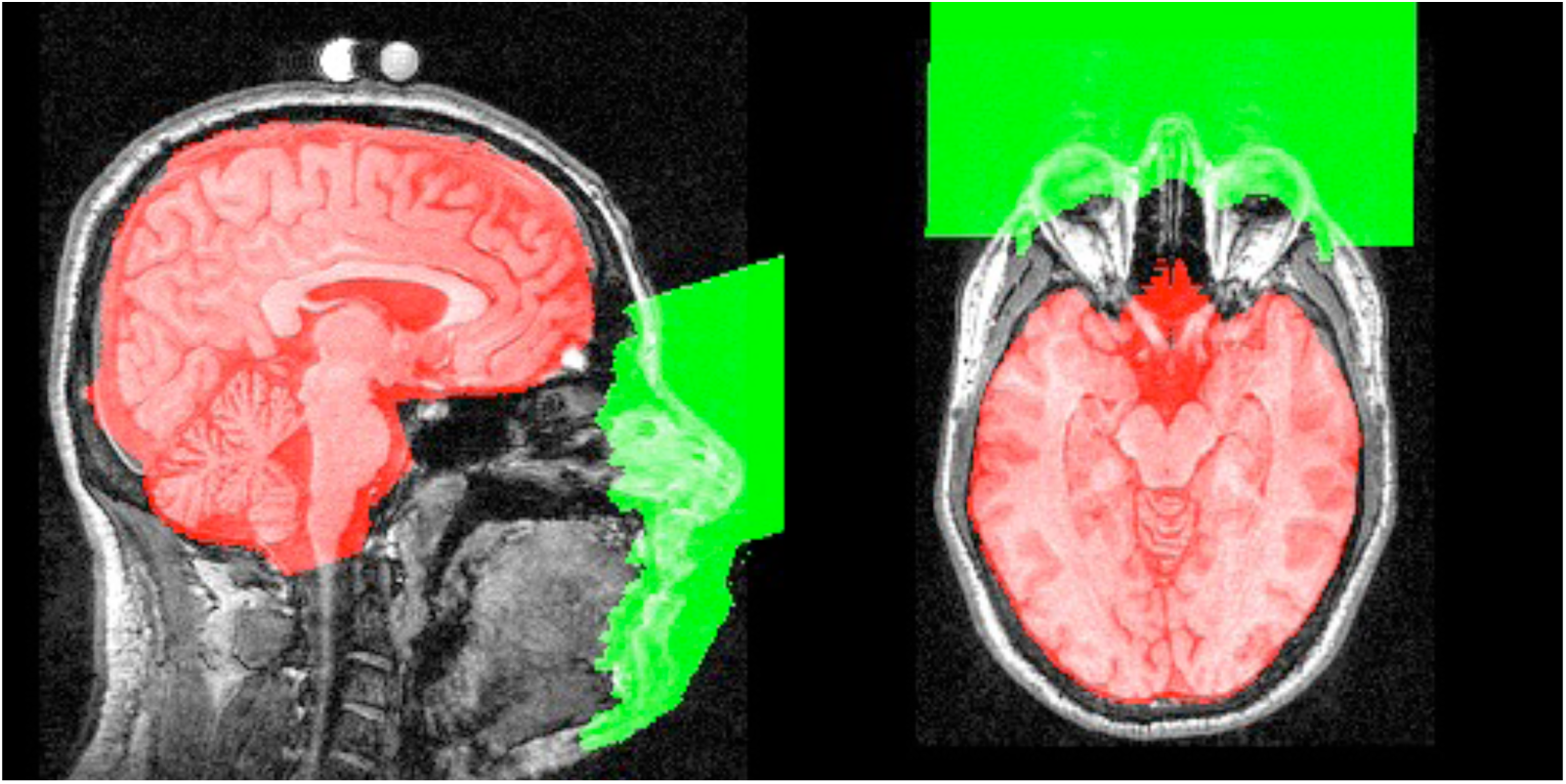
Defacing masks: red - brain mask, green - face mask.

To deface the subject a random smoothed vector field 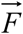 characterized by standard deviation of each component *σ* and maximum amplitude *A* is used. It can be produced by convolving a random vector field with a gaussian kernel and normalizing the maximum amplitude of the resulting field.

To enforce a smooth transition from the undistorted part of the skull to the distorted part, a smoothing face mask *M* _*fs*_ is used, it is produced from the face mask *M* _*f*_ by calculating the chamfer distance function and thresholding it by a given smoothing distance *S* and normalizing the result in the range [0: 1]. The smoothing face mask *M*_*fs*_ is then multiplied with 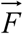 element-wise, producing the resulting defacing field 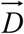. The defacing vector field is then transformed into native space for each modality by applying the inverse transformation: 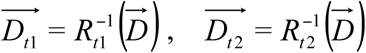 and 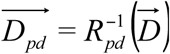. Scans in T1w,T2w and PDw modalities in native space are then resampled using the defacing vector fields that results in pertubations being only applied the facial tissues. Figure 2.

**Figure 2.**
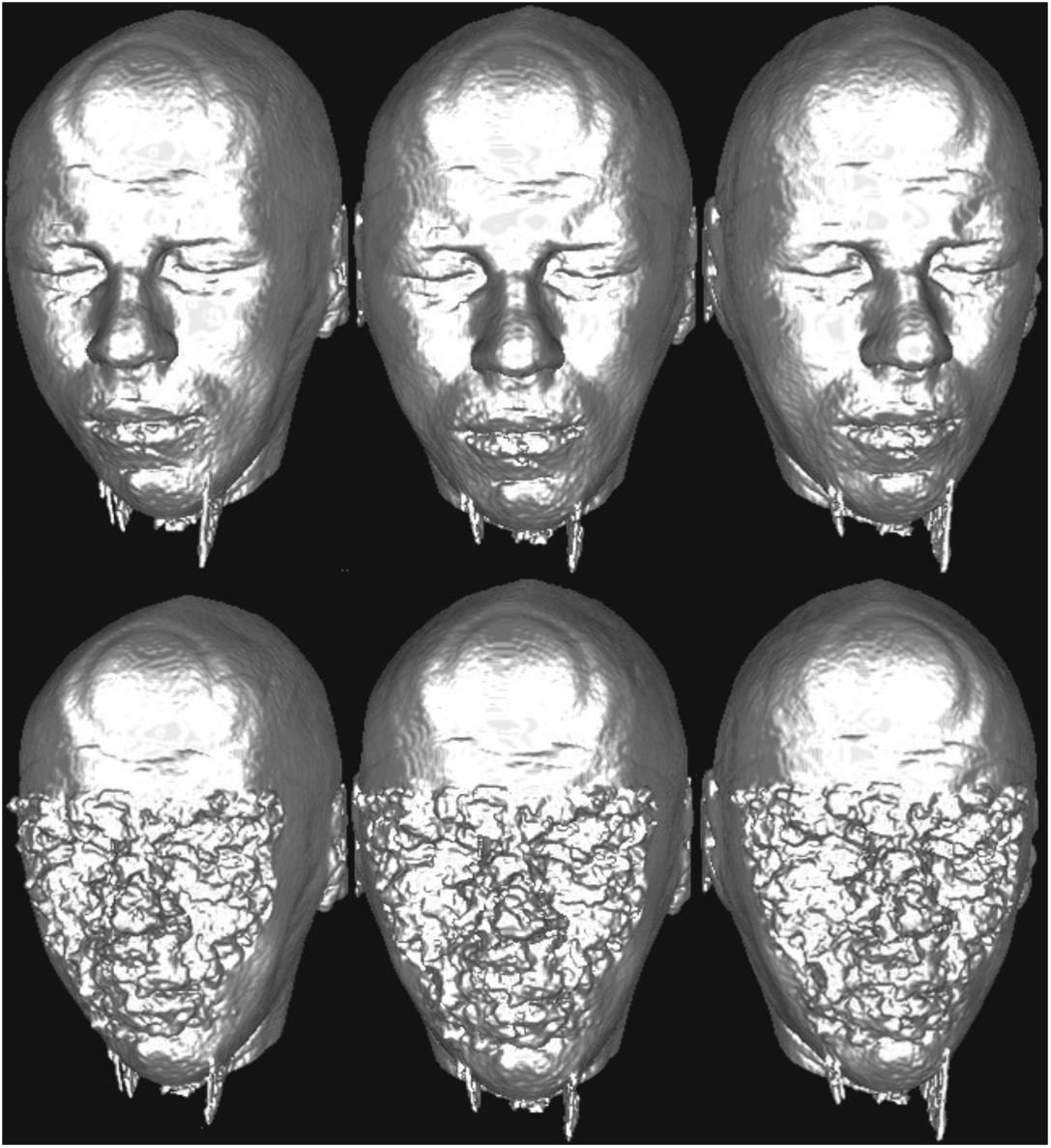
Rendering of a face before (top) and after (bottom) defacing.

### 2.2 Validation

To demonstrate that the proposed defacing procedure does not significantly affect subsequent data processing a validation using 149 of the ICBM database was performed. First, the effect on automated stereotaxic registration was measured by comparing the linear registration transformation of the original and defaced scans. For each voxel *v* withing the brain of a subject we estimate the vector 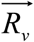 which brings this voxel into stereotactic space and find the maximum difference within the brain between the normal and defaced subject 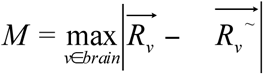, Figure 3 shows the distribution of this distance between subjects:

**Figure 3.**
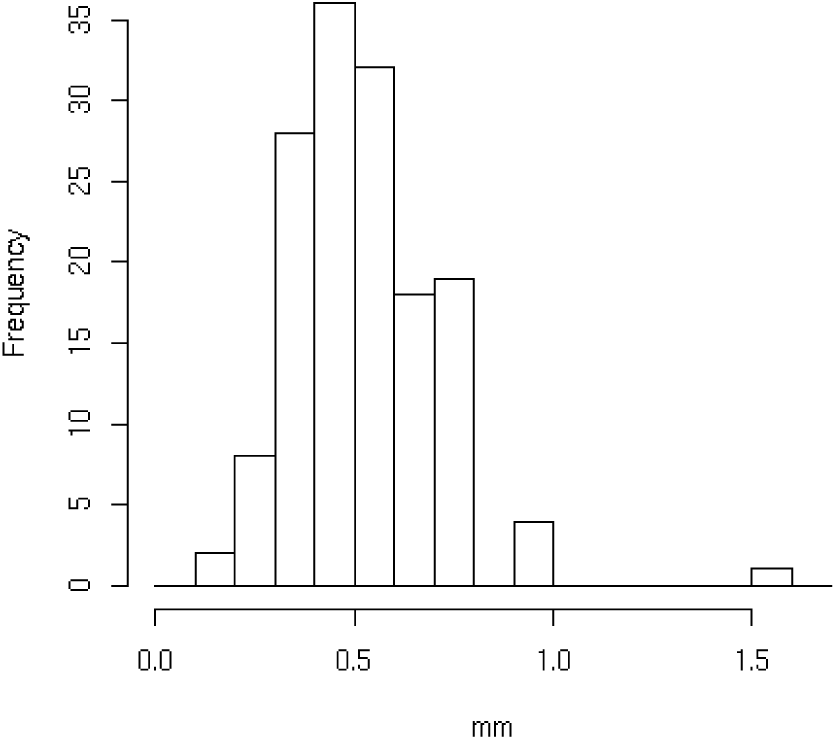
Maximum distance after registration withing brain.

To verify that tissue classification is not affected by defacing we use the corrected *κ* measure [8] between labeled volumes produced from the normal and defaced scan, figure 4. Corrected *κ* [15] is defined as: *P*_*o*_ - observed percentage of agreement, *P*_*c*_ -percent agreement that would occur by chance alone: 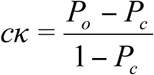

**Figure 4.**
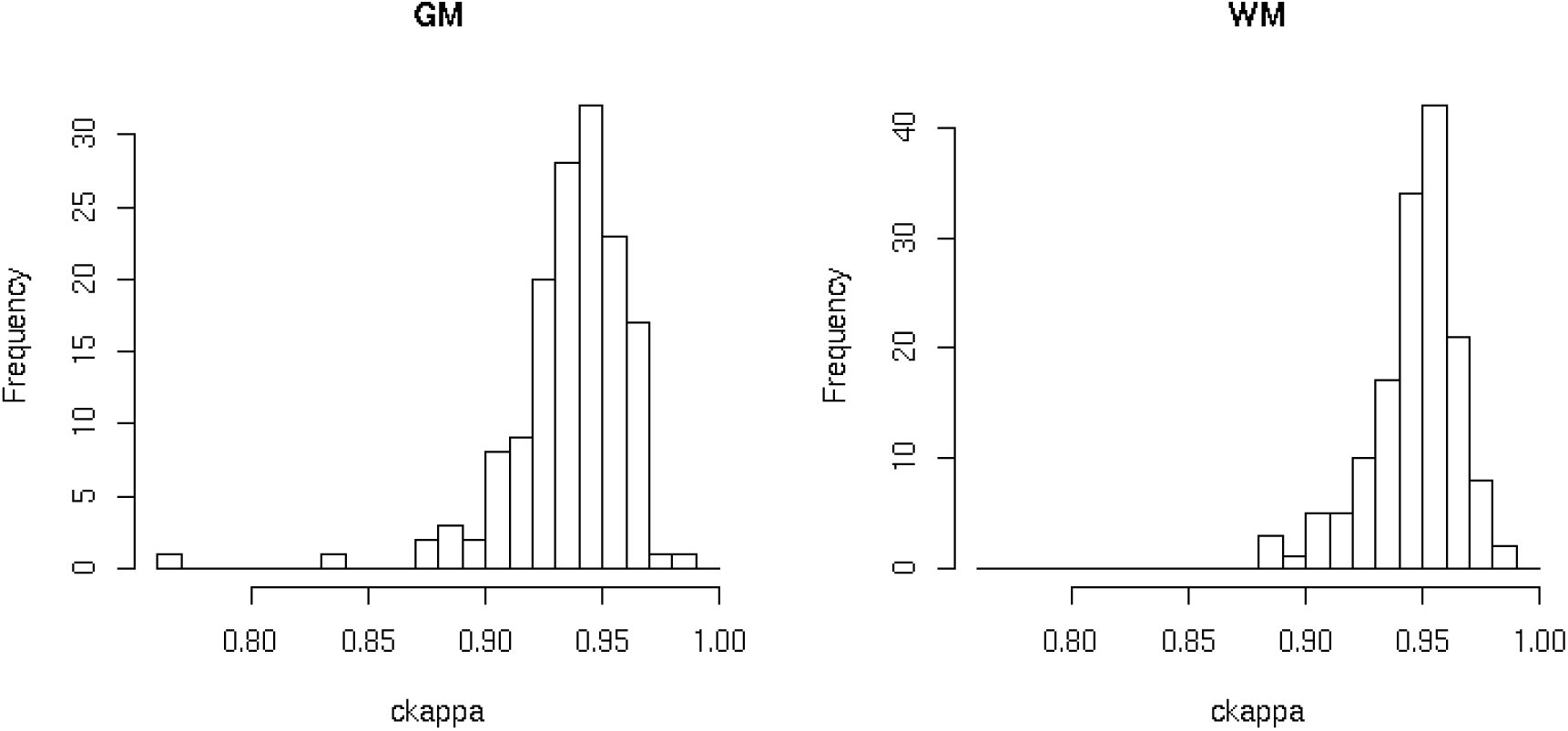
Corrected kappa scores for Gray Matter (GM) and White Matter (WM).

## 3 Discussion

This paper has demonstrated and validated a method that changes the appearance of a subjects volume rendered MRI data beyond recognition and in doing this does not significantly alter features of the image to the point that subsequent processing is significantly altered. The algorithm that has been described has been demonstrated to work on T1w, T2w and PDw data but clearly is extensible to any other modality of data so long as a inital stereotaxic registration transformation can be calculated. The degree of defacing is controllable by two parameters, the FWHM of the blurring kernel that is applied to the random field and the amplitude of the random field. In the examples above a FWHM of 10.0mm and amplitude of 5mm was used. These parameters can be changed for differing head sizes or data sources but these values will suffice for most purposes.

The described method takes approximately 1 minute per scan on medium level hardware available at the time of writing this paper (2 GHz PC) for a 256×256×256 image. As such, this method is very applicable to large imaging databases such as the NIHPD study and is completely automated once the inital probabilistic mask has been determined. In fact, this method was successfully applied to the NIHPD database, and the resulting face rendering were manually inspected by an expert and results were found acceptible. The method is implemented as part of the *standard_pipeline.pl*, see https://github.com/BIC-MNI/bic-pipelines

